# Evolving building blocks of rhythm: How human cognition creates music via cultural transmission

**DOI:** 10.1101/198390

**Authors:** Andrea Ravignani, Bill Thompson, Thomas Grossi, Tania Delgado, Simon Kirby

## Abstract

Musical rhythm, in all its cross-cultural diversity, exhibits several commonalities across world cultures. Traditionally, music research has been split in two fields. Some scientists focused on musicality, namely the human biocognitive predispositions for music, with an emphasis on cross-cultural similarities. Other scholars investigated music, seen as cultural product, focusing on the large variation in world musical cultures. Recent experiments found deep connections between music and musicality, reconciling these opposing views. Here we address the question of how individual cognitive biases affect the process of cultural evolution of music. Data from two experiments is analyzed using two different, complementary techniques. In the experiments, participants hear drumming patterns and imitate them. These patterns are then given to the same or another participant to imitate. The structure of these - initially random - patterns is tracked down to later experimental ‘generations’. Frequentist statistics show how participants’ biases are amplified by cultural transmission, making drumming patterns more structured. Structure is achieved faster than in transmission within, rather than between, participants. A Bayesian model approximates the motif structures participants learned and created. Overall, our data and model show that individual biases for musicality play a central role in shaping cultural transmission of musical rhythm.

## Introduction

Empirical musicologists distinguish music from musicality ^1^. Music is a cultural product, while musicality is the human biological machinery used to produce and process music. This is a helpful distinction, especially when asking evolutionary questions. To a first approximation, how music changes over time seems orthogonal to the evolution of the biocognitive apparatus underlying music-making. Recent experiments, however, have shown that music and musicality are intimately connected ^2-6^. Focussing on rhythm, individual participants in ^4^ were asked to imitate snare-drum sequences on an electronic drum set to the best of their abilities. All participants were non-musicians, and were not told where the patterns came from. A ‘first generation’ of participants was given computer-generated drum patterns, featuring random beat intensity and duration. Successive ‘generations’, however, were asked to imitate the output of the previous participant in the experiment (Figure 1A, top ‘chain’ of participants). Unbeknownst to each participant, patterns included all the errors and imperfections introduced in the previous generation. In this way, the process of cultural transmission of rhythmic patterns was recreated in the lab ^2, 7, 8^. Over time, these sequences became more structured and easier to imitate. In addition, rhythmic sequences converged towards all rhythmic universal features which characterize music around the world ^9, 10^. This experiment, enriched by similar findings in non-musical domains ^11-16^, suggests a link among cognition, biology, and culture in human music ^17, 18^. In particular, features present in almost all musical traditions around the world emerge through basic bio-cognitive biases amplified by the process of cultural transmission. This experiment also raised a number of additional questions. In particular, are these bio-cognitive biases specific to: (1) humans, (2) adults, (3) Westerners, (4) a cognitive domain or modality, and (5) particular individuals? The question of individual-specificity has broad implications no matter its answer. In one scenario, if all individuals had similar biases, music would emerge as a result of strengthening few, human-widespread tendencies towards musicality (akin to a vote by consensus). In another alternative scenario, if all individuals had different biases, music would emerge as a result of the interplay among individual-specific tendencies towards musicality, amplifying or averaging each other out (similarly to mixing paint of different colours). In other words, do regularities in musical structure result (a) from distributed, but largely homogeneous, weakly-biased processing or instead (b) from specific individuals who impose idiosyncratic structural regularity ^19^?

**Figure 1:**
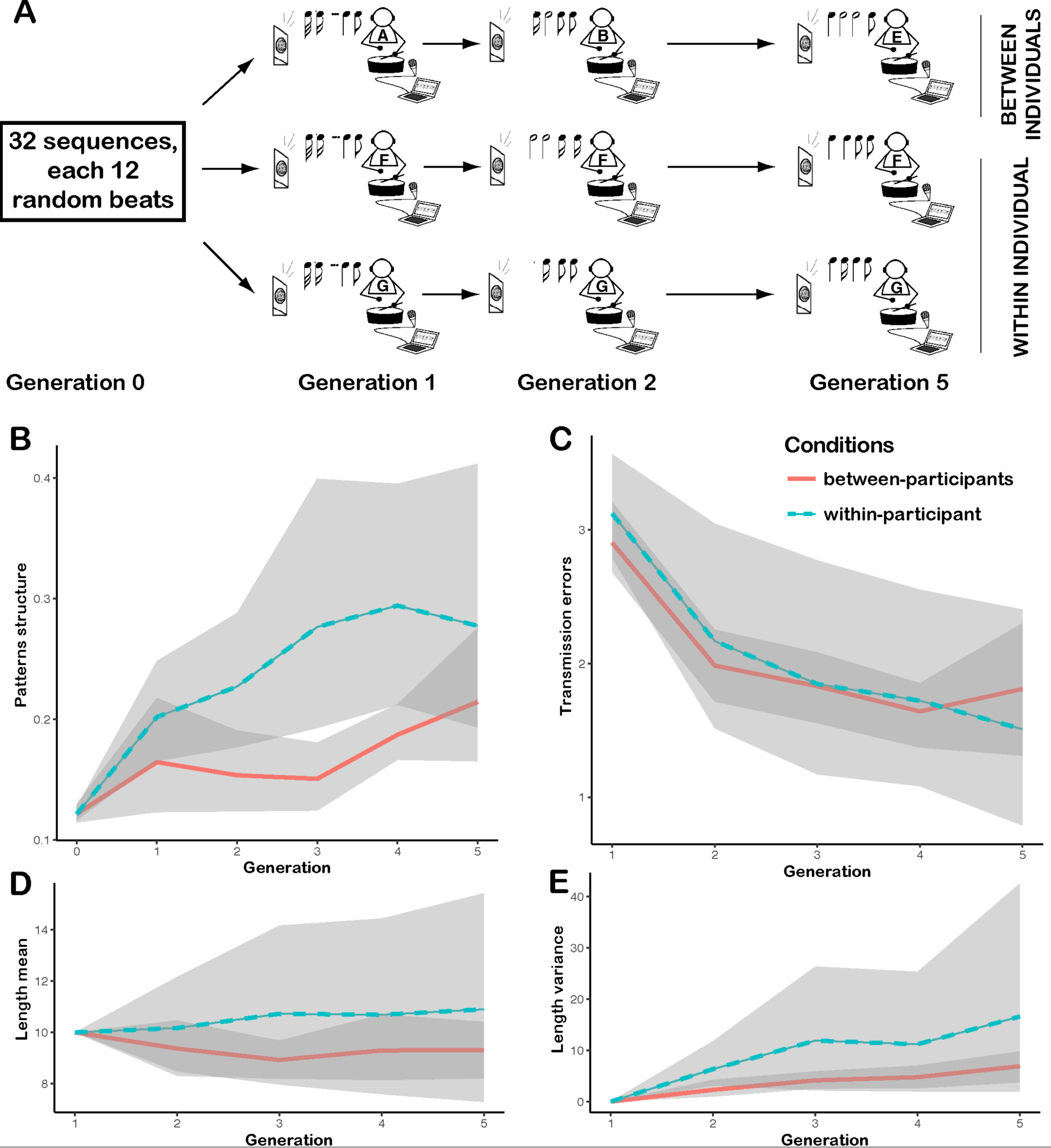
Experimental design and summary statistics tracking the evolution of patterns in the first 6 generations of both experiments. (A) The experimental design follows a transmission chain paradigm: the output of one ‘experimental generation’ constitutes the input of the next generation. Generation 0 consists of computer-generated drumming patterns, where drum hits have random velocity and inter-onset intervals (IOIs). Each generation 0 pattern is individually heard and imitated by a Generation 1 participant. The resulting imitated patterns can be given to the same participant to imitate once again (bottom two rows of A, within-individual design), or to a different participant (top row of A, between-individuals design). The procedure is repeated over generations (left to right), and additional chains (not shown). (B) Increase in structural complexity quantified over generations using a modified measure of entropy. (C) Decrease in imitation error between adjacent generations, quantified using a modified Levenshtein distance. (D) Pattern length across generations and chains. (E) Variance in pattern length across generations and chains. (B-E) Shaded areas depict bootstrapped 95% confidence intervals, lines connect the mean value for each generation and experiment type across chains.

In this paper, we start tackling the issue of how individual biases might affect the process of cultural evolution of music. In particular, we ask whether similarity in biases within a transmission chain can speed up the process of cultural evolution. We address this question using experimental manipulations and Bayesian modeling techniques. First, we replicate the original experiment in ^4^ with one variation. In the original experiment, a set of drumming patterns was transmitted across generations of participants in between-participants design (top row of Figure 1A). In contrast, the current experimental design features a within-participant structure (bottom two rows of Figure 1A) ^3, 19-21^. Each participant takes part in multiple rounds, instead of one round per participant as in the previous experiment. Second, we analyze and compare the data from the two experiments using standard inferential statistics. Third, we introduce a probabilistic model for the latent structures underpinning rhythmic sequences, alongside a psychologically plausible algorithm for inferring these structures. This allows us to obtain approximate structural descriptions of rhythmic patterns across conditions and generations, and to explore how these structures are used and re-used. Whereas previous models have focused on inferring cognitive biases for integer ratio rhythmic categories from experimental data ^3^, our model focuses on approximating the process through which individuals combine rhythmic categories into predictable motif-like sequences.

A general prediction is that the new within-participants “chains” will likely produce data *qualitatively* comparable to the original between-participant design. In particular, patterns should increase in structure, and become easier to learn. If this holds, self-learning will be shown as an effective method for uncovering musical biases in participants. *Quantitatively*, however, the two experimental designs might show different behaviours. In particular, if participants have individual-specific biases, they will likely impose an idiosyncratic structure every generation. This, in turn, will make structure emerge faster in the within-participant chains, and slower in between-participants chains (some innovations will cancel out). If this is the case, we should observe a significant difference between the variables measuring the evolution of structure in the two experiments. If, instead, participants have homogeneous biases towards rhythmic structures, the evolution of chains will not be affected by experimentally substituting many participants for one repeated participant. In this case, the two experimental designs will produce similar data, hence no measurable difference between key variables in the two experiments.

### Experimental Methods

Data from six experimental chains (30 participants) from a previous experiment were reanalyzed (for details see ^4^). In brief, six different sets of 32 sequences of 12 random beats were given to ‘first generation’ participants to imitate. First generation output became second generation input, and so on (see Figure 1). In addition, new data was collected from 12 experimental chains (12 participants) in conditions comparable to the previous experiment ^4^, with two key differences (for details see Supplement). First, each of the six different sets of 32 sequences of 12 random beats used in the first experiment was given to two first-generation participants in this new experiment (as opposed to one first-generation participant in the previous experiment, see Figure 1). Second, the new experiment featured within-individual transmission, so that the same participant listened and imitated their own drum patterns over 5 experimental generations. In both experiments, the participant was unaware that she would imitate her own, or someone else’s, previous pattern

### Frequentist statistics: Results and discussion

The metrics tracking structure and imitation error behave similarly over generations (Figure 1B, 1C), confirming our qualitative prediction. We used ANOVAs to test whether the ‘generation’ and ‘transmission type’ (i.e., between or within-individual) could account for a possible increase in structure ^4, 22, 23^ (Figure 1B) and decrease in imitation error ^4, 24^ (Figure 1C). A stepwise model selection suggested that both generation and transmission type should be entered in the ANOVA as predictors of structure (minimizing Akaike Information Criterion). Both variables were significant predictors of structure (transmission type: F=7.4, p<.01; generation: F=14.5, p<.001). Another stepwise model selection suggested that only generation should be entered in the ANOVA as predictor of imitation error. Generation was a significant predictor of imitation error (F=14.8, p<.001).

We used further ANOVAs to test that these differences were not due to superficial features. We calculated the length of each drumming pattern (in the ‘r space’, see Figure 2), and computed mean and variance length for each set of 32 patterns. A stepwise model selection suggested that neither generation nor transmission type should be entered in the ANOVA as mean pattern length. Another stepwise model selection suggested that only generation should be entered in the ANOVA as predictor of variance in pattern length. Generation was a significant predictor of variance in pattern length (F=4.0, p<.05). In other words, while participants vary the number of beats produced within an experimental session across generations, this does not appear affected by the transmission type. Crucially, transmission type affects neither mean nor variance in pattern length, suggesting that structural variability across experimental groups is not attributable to simple differences in pattern length.

**Figure 2:**
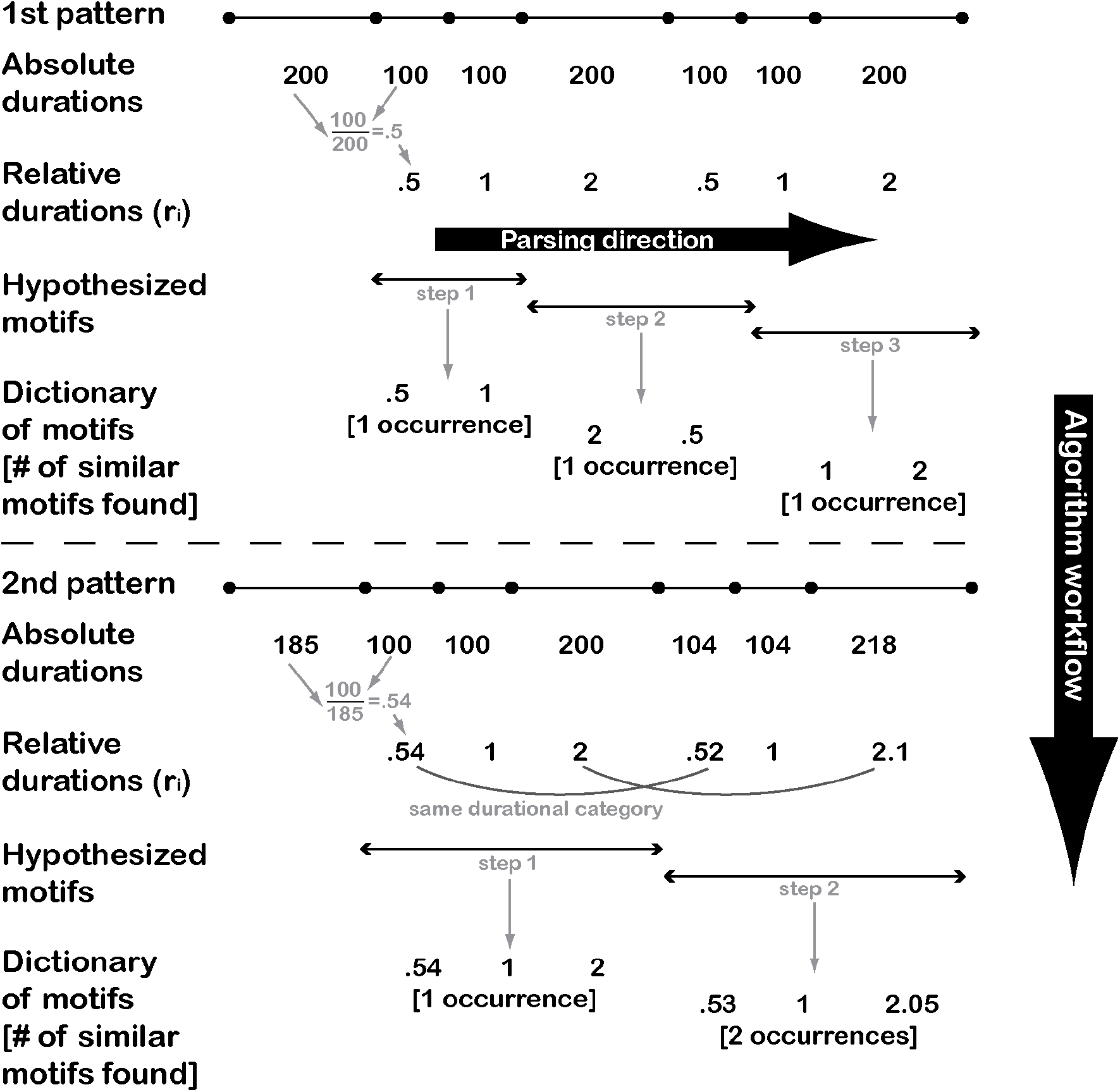
Latent variables Bayesian model: Sketch of how the algorithm processes two drumming patterns (workflow proceeds from top to bottom, and from left to right). A drumming pattern (first pattern, topmost row) can be conceptualized as a series of duration marked by drum events (lines broken by circles). The (absolute) durations between drum events are represented as a vector of IOIs. By taking the ratios between adjacent IOIs, one obtains a vector of relative durations r=(r_1_,…,r_n-2_). All IOI sequences with the same r vector have the same rhythmic pattern up to a tempo multiplicative constant. The algorithm generalizes first over r_i_ categories (e.g. in the first pattern .5 and .5 belong to the same category) and then assigns every hypothesized motif either to its prototypical category or to a new category. When the participant hears a new pattern (second pattern) with more variability, ratios such as .54 and .52 might be assigned to the same r_i_ category. Likewise, the algorithm randomly attempts to be ‘greedier’, probing the existence of motifs of length 3 or above, hence finding that sequences like (.54, 1, 2) and (.52, 1, 2.1) belong to the same category.

Together, these results suggest that both structure and learnability change over time. Although transmission type does not affect learnability, it does affect the amount of structure: a within-participant design results in higher levels of structure. This provides preliminary support for the ‘idiosyncrasy of biases’ hypothesis over the ‘homogeneity of biases’ hypothesis. These inferential statistics suggest that repeated idiosyncrasies enhance the emergence of structure (within-participant transmission type). Conversely, new minds introduce more variance, slowing down the emergence of structure (between-participants transmission type). The inferential statistics, however, are not able to unveil the structures participants infer. In the next section, we outline a mathematical model as an approximation to what participants perceive and learn.

### Computational model: Methods

Data from both experiments were further analyzed using a computational model (see Supplement). Our model formalises the idea that participants may decompose individual drum patterns into sequences of motifs that can be reused across patterns. Given a drum pattern, the model attempts to infer boundaries between latent motifs, and to categorise these motifs into coherent groups based on prototypes. Our approach takes inspiration from two related fields. First, since the inferential task is essentially one of joint segmentation and clustering, we can adapt techniques from the machine learning and speech technology literature ^25^ to specify a probabilistic model of underlying latent variables (motifs and their boundaries). Second, the literature on human statistical learning provides a psychologically plausible algorithm that makes guesses about these unobserved variables ^26^.

Our model formalises two levels of structure within a pattern, by grouping adjacent interval ratios into a small number of Gaussian categories via Bayesian inference ^a^, and by using sequences of these inferred categories to construct an inventory of motifs that can be reused. Our posterior approximation algorithm learns this structure ‘on the fly’ in a probabilistic, sequentially dependent, psychologically plausible fashion: as new elements of a drumming sequence are perceived, the backwards transition probabilities (BTPs) between interval ratio categories are estimated ^23^, and used to hypothesise boundaries between motifs. The model tries to assign any hypothesised motif to an existing category of motifs (Figure 2, top), and creates a new motif category whenever this fails (Figure 2, bottom).

A candidate subsequence has to go thorugh two criteria to be considered a motif: (1) how often has the model seen this particular subsequence of categories? (2) how often have the preceding and current element been found together (BTPs)? If the same sequence has been seen before, this provides evidence for the current subsequence to be another occurrence of this motif type. If the first element of a subsequence and the previous element rarely co-occur, this provides evidence that the two elements belong to different motifs, implying a boundary. The model includes parameters which influence, for example, how willing the learner is to invent new motif categories. These parameters and all other details of the model are described in detail in the supplement to this article. In the analyses we present here, we set these parameters such that the model is weakly biased to prefer re-using existing motifs, but able to invent new categories whenever the data dictate. Because the model includes a free parameter that determines this balance of re-use and invention, this assumption could be straightforwardly revisited in future analyses. Crucially, we fix parameters to be identical across analyses of both experimental datasets, and examine how the structures inferred by the model vary by experimental condition (rather than examine the specific structures inferred, which can be sensitive to model parameterization).

### Bayesian model: Results and discussion

We ran the model through both experimental datasets 10 times, which (insofar as our model is a psychologically plausible theory of participants), provides an approximation to the representation of patterns induced by participants. We examined two principal measures of structure: 1) the number of unique motifs discovered by the model at each generation of each chain, and 2) the number of patterns in which each attested motif was discovered at each generation of each chain. Together these measures allow us to quantify the evolution of structural regularity across a set of rhythms, over generations, as a function of how participants learn and the data they produced.

In both experimental conditions, the number of unique motifs attested within a generation decreased over generations (Figure 3, top row). In line with our inferential statistics, this suggests an increased degree of re-use of prototypical building blocks over generations, and that these building blocks are discoverable by a simple algorithm making local, sequential decisions. The regression slope plotted in Figure 3 show that this happens faster in the within-subjects chains, and that the fifth, and final, generation of within-subjects chains re-use motifs to a slightly greater extent than the equivalent generation of between-subjects chains.

**Figure 3:**
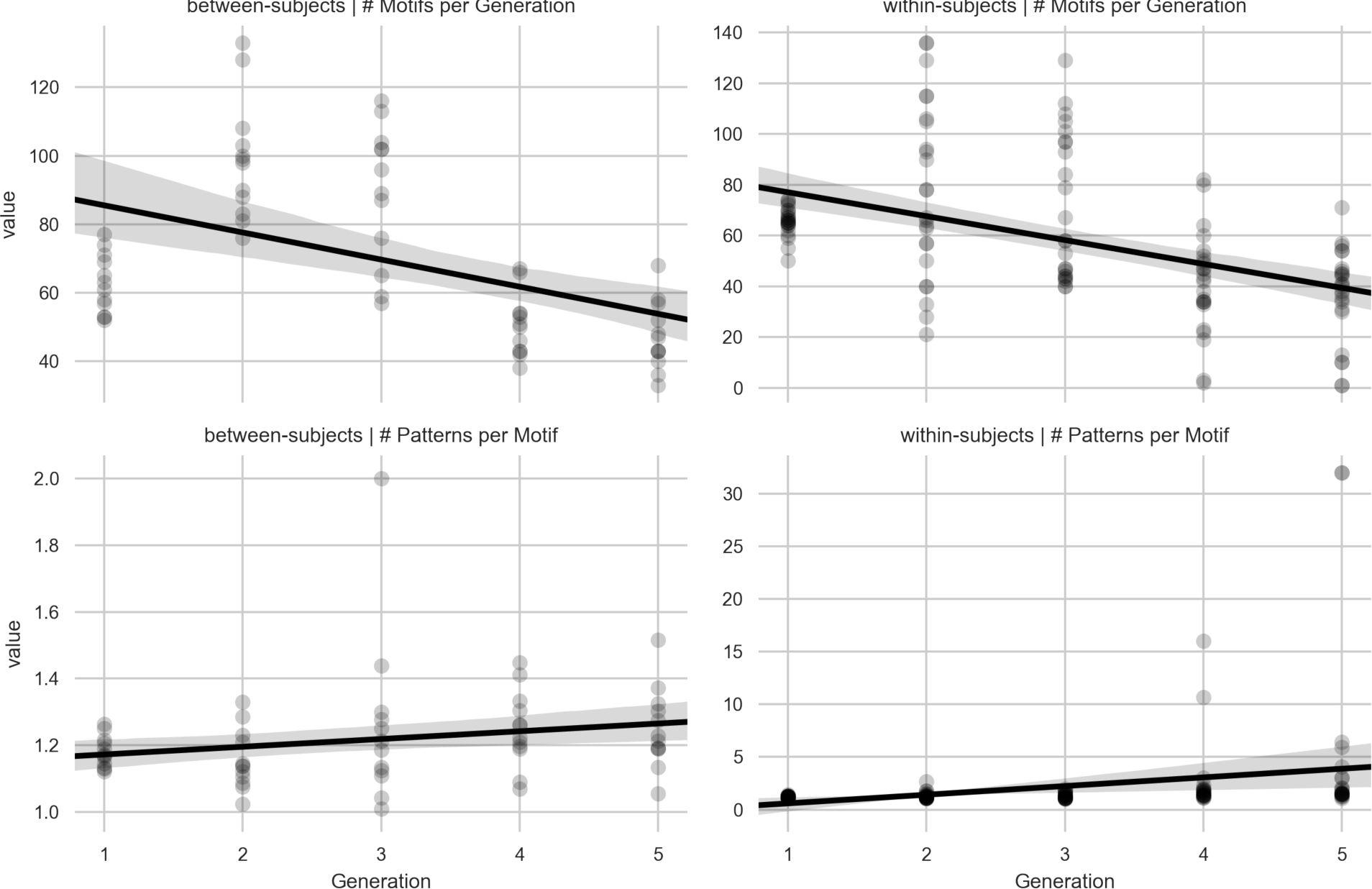
The number of motifs inferred by the model (averaged over 10 independent simulations) at each generation (top row) in the within-subjects (right) and between-subjects (left) chains, and the number of independent patterns in which each motif evidenced at least once in a generation was identified in that generation (bottom row). Points show these quantities for individual motifs (simulated ten times); lines show regression slopes.

Our analyses also suggest (Figure 3, second row) that in within-subjects chains, each attested motif tends to be present in an increasing number of patterns over generations, suggesting participants are entrenching the motifs they have invented. This pattern is also visible in the between-subjects chains, but to a lesser extent. While these chains do evidence increasing reuse of motifs, they do not appear to evidence the same degree of entrenchment on a small set of widely re-used motifs as is suggested in the within-subjects results. We interpret this as a sign that (1) both experimental conditions lead to an increase in structure, but (2) the within-subjects condition allows the idiosyncrasies of individual minds to repeatedly bias the distribution of structures in a chain-specific way that is less probable when new learners are forced to re-interpret the structures invented by previous individuals.

Finally, we also computed the distribution of interval ratios at each generation (Figure 4). Both experimental conditions evidence a sharp transition from unstructured initial distributions (generation 0, top row) to highly structured categories of intervals. Interestingly, while the tri-modal distribution with peaks near interval ratios is clear in the between-subjects data (as previously reported ^4^), the within-subjects chains converged on an approximately two-way category distribution with peaks at 1 / 2 and 1 ^3, 27^.

**Figure 4:**
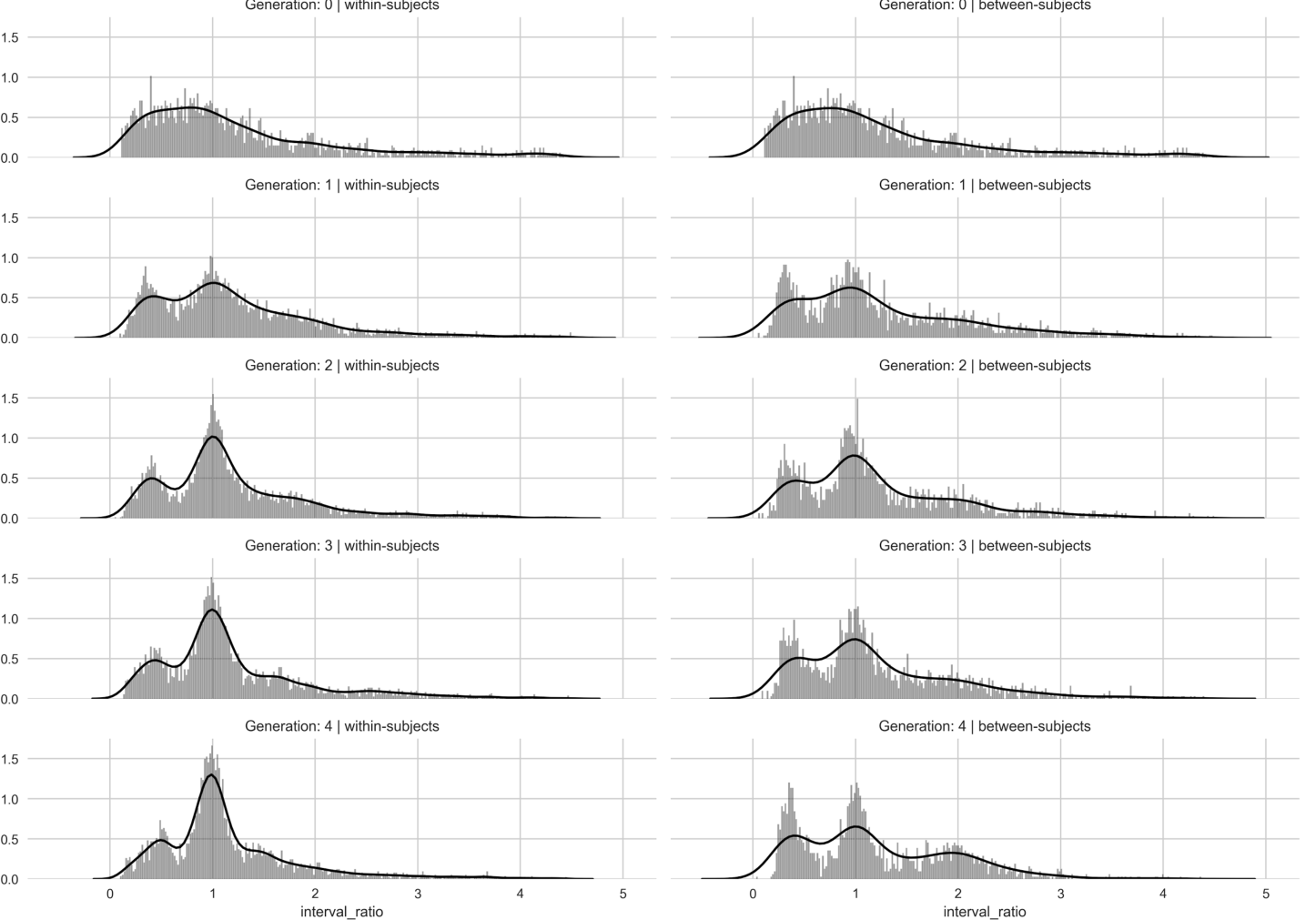
Normalised histograms for the distribution of interval ratios at each generation in the between-subjects (right) and within-subjects (left) chains. Lines show kernel density estimates of these distributions.

### General discussion and conclusions

In this paper, we investigate the role of individual cognitive biases in the creation of rhythmic patterns. We present two experiments in non-musicians, analyzed using two complementary techniques. We show, in a musical imitation task, that rhythmic regularities emerge via simulated cultural transmission. Similar regularities emerge when participants are asked to imitate their own or other participants’ patterns. However, this design affects the amount of regularities emerging: When participants imitate their own previous productions, convergence is faster. We suggest this is due the presence of weak idiosyncratic biases. When transmission occurs between participants, idiosyncratic biases partially cancel each other out. Instead, when transmission occurs within a participant, biases reinforce each other. Our conclusions partly contrast with computer simulations done for language, where in heterogenous populations a few oulier agents distort the signal transmited in an otherwise heterogeneous population ^19^.

The study has several limitations, which could be improved upon with future work. First, we adapted and designed the model to have cognitive plausibility and match experimental conditions as closely as possible. However, our Bayesian model still neglects several findings from rhythm perception and production ^28, 29^, and should be understood as a first approximation. The model should be refined to include additional empirical findings, becoming as psychologically and neurally plausible as possible. Moreover, while we focused on psychological plausibility by implementing a sequential posterior approximation algorithm as our model of the learner, future analyses might instead focus on inferring the best possible structural descriptions, by implementing more computationally intensive posterior sampling procedures. Second, one dimension we completely discard in this work is the intensity information of each beat. Velocity and accents are an integral part of rhythm, so future extensions of this work should go beyond pure durational information. Third, the Bayesian model includes a number of parameters which undoubtedly influence our results. Future modelling efforts might profitably aim find empirical motivation for these parameters, or derive simpler models with fewer parameters. Ours is only a first attempt, that is why we are happy to share our data and scripts with interested researchers who would like to perform modifications. Fourth, given a restricted set of assumptions, we have predicted the number and distribution of rhythmic motifs inferred by participants. Ideally, whether participants actually acquire similar motif-like substructures will need to be tested experimentally by asking participants to classify motifs, and check how closely these decisions align with the model’s prediction. Fifth, while one would expect less of a difference if priors were homogenous, the design of the two experiments differs along one additional dimension. In fact, in the within participant experiment, participants can potentially carry memory over from previous generations. If this was actually happening, one would actually predict a slower evolution of structure in the within participant condition (since the effect of the prior would be relatively weakened as there’s a drag from the increasing pile of data). The fact that this does not happen may suggest that - at least in this design - participant memory is not a crucial factor. In fact, subjects in the within participant condition were not told they were listening to their own data. That, plus the sheer number and length of sequences, and the participants' lack of musical training should explain why memory isn't as strong a factor. Future research will be faced with disentangling biases’ homogeneity from memory effect. Participants’ grouping based on their electroencephalographic signature might be a viable solution ^30^.

We believe this research makes a contribution to a number of disciplines. Over the past century, a deep divide has separated cultural anthropology and music psychology ^31^: our experiments try to bridge this divide with a design that takes into account both the cultural medium and human bio-psychological features. Likewise, our purely behavioural experiments could be combined with neuroimaging or electrophysiology techniques to tap into the neurobiological bases of human biases for musical rhythm. Within the interdisciplinary field of cultural evolution, we show how within-participants transmission speeds up the process of convergence. Quantitative models abound in music information retrieval, but are still scarce in music cognition: Here we adapted some recent computational techniques to the interpretation of human data. In brief, we hope that our paper will spur a tighter integration of modelling and empirical research in the study of the psychology and neuroscience of music.

## Acknowledgments

AR and BT are grateful to Marieke Woensdregt and Carmen Saldaña Gascón for hospitality and support during the development of this research. Andrea Ravignani has received funding from the European Union’s Horizon 2020 research and innovation programme under the Marie Skłodowska-Curie grant agreement No 665501 with the research Foundation Flanders (FWO) (Pegasus^2^ Marie Curie fellowship 12N5517N), and a visiting fellowship in Language Evolution from the Max Planck Society.

Authors contributions: AR and SK conceived the research, AR, TG, TD and SK designed the experiments, TG and TD performed the experiments, all authors analyzed the data, AR, BT and SK conceived the model, BT implemented the model, AR and BT wrote the manuscript, all authors edited and approved the manuscript for publication.

## Conflicts of interest

The authors declare no conflicts of interest.

**Figure legends**: Should be provided at the end of the manuscript.

**Footnotes and Endnotes**: (Use lower-case italic letters in superscript.)

^a^Each of the 32 imitated pattern can be described as a time series of inter-onset intervals, i.e. the time between adjacent drum hits IOI_1_, IOI_2_,…, IOI_n_. As in the original study, to account for possible tempo drift within and across patterns, we use the ratio between adjacent beats, i.e. r_i_=IOI_i+1_/IOI_i_. Hence each pattern of *n* hits can be represented as a time series r_1_,…,r_n-2_. The computational model proceeds by first clustering data points into rhythmic categories. We assume that participant do not perceive and represent the absolute magnitude of the r_i_ data points, but potentially reduce the variation among data points by assigning each to a rhythmic category.

## 1. Experimental Methods

### 1.1. Participants

12 participants (7 males) were recruited using the University of Edinburgh’s graduate employment service (mean age 24.25). The sample size was established a priori and based on number of experimental chains (instead of participants) for comparability with the previous study. All participants were paid £30. Conditions for participation were the lack of any formal musical education, to any degree.

### 1.2. Materials

Stimuli for the experiment were sequences of midi drum beats, stored as lists of numbers. Each drum beat was defined by two values, velocity (on a scale of 0-127), representing hit strength, and interonset-interval (IOI), representing time in seconds between beats. To play a sequence to the participant, the experiment code transformed the velocity values into midi output (midi note 40, snare drum), separated by intervals determined by the IOI values. After all beats had been played, there would be 1.5 seconds of pause, and a single midi note of 32 (cymbal) would signal to the participant that the sequence had ended. Midi output was converted into audio by the software Garageband, set to its percussion channel (standard rock kit).

The training stimuli for each round were 32 snare drum sequences. Presentation of these 32 basic sequences was repeated twice, each time randomizing their order. After a sequence was played, the participant would copy it on a midi drum set. A custom Python script collected values of velocity and IOI from the participant’s input and stored them. The 32 basic training sequences for the first generation of each chain were identical to the 32 basic training sequences used in our previous experiment (Ravignani et al., 2016). Briefly, first-generation sequences comprised 12 beats, whose values of velocity and IOI had been randomly generated (each IOI was randomly sampled from a uniform distribution between 100 and 1,000 milliseconds). For all other rounds of the experiment, training sequences were built using the last 32 output sequences of the participant’s previous round, again produced twice in random order. Sequence length was left free to vary, with minimum length being 1 beat.

The experiment was run on a MacBook Air (OS 10.8.5), which received midi input from a midi drum-kit (Alesis SamplePad). Drum-kit and laptop were connected via an USB-midi external interface (Roland Duo-Capture Ex). Sound were played over headphones connected to the USB-midi interface, whose volume the participant was free to adjust at the beginning of the experiment. The participant received a single standard wooden drumstick.

The script used to collect data was a modified version of the Python script in (Ravignani et al., 2016). The only difference consists in the new script using the (better documented) mido and time modules instead of the rtmidi module. This alteration did not lead to a noticeable difference when replaying the sequences.

### 1.3. Design

This study aimed at replicating the six diffusion chains in (Ravignani et al., 2016). Each of the original chain was used here as generation 0 for two participants. This lead to a total of 12 chains (versus 6 in the previous study). All rounds of a single experiment were run on the same day, to control for potential beneficial effects of sleep on learning (Maquet, 2001). In an effort to reduce the overall duration of the experiment, and considering the little variation in the original chains after generation 5, each chain consisted of 6 generations: one computer-generated generation 0, and 5 experimental generations. Chains were named after the shared input with original chains, with chains 1 1 and 1 2 being replications of chain 1. Participants underwent five rounds of 20-25 minutes each, separated by four 15-minute breaks. Each experiment lasted approximately 3 hours.

### 1.4. Procedure

Participants were given consent forms, which included, among others, the instructions: “You are about to participate in a study which involves listening to sequences of drum beats and immediately repeating them on the electronic drum kit provided by the experimenter, to the best of your abilities. You will be shown how to use the kit before starting. The experiment is split into five rounds, each lasting approximately 30 minutes and followed by a break of approximately 15 minutes. Your session should last for up to 3.5 hours. You will be given full instructions shortly and will be able to ask any questions you may have.”

After signing the consent forms, participants were given headphones and a drumstick. Apart from the written information in the form, participants were debriefed by the experimenter. They were told they would listen to sequences comprised of “certain number of drum beats”, and they would know the sequence had ended when they heard a cymbal sound. They were given a few moments to test the pads and recognise that the top-right pad produced the cymbal noise, and were told that that pad should only be used to signal the end of their own sequence. Once they were satisfied with the range of sounds they could produce, participants were told a round would last approximately 25 minutes, and there would be five of these rounds, separated by four breaks of up to 15 minutes. Participants were unaware whether they would listen to stimuli produced by a computer, a previous learner, or themselves at a previous time.

Before starting, participants were given three recommendations. Firstly, they were not expected to replicate the sequences perfectly, but they were expected to “make an honest effort to replicate them to the best of” their abilities. Secondly, if they heard a different drum sequence in their headphones while still replicating the previous one, it meant they had hit the cymbal pad by accident, and the code had given them the next sequence: they should therefore stop, listen to the new sequence and replicate that one once cued by the cymbal sound. Thirdly, the experimenter would be just outside the experimental room if they were feeling ill or faint or something important had come up, but barring those events, they should only come out once they failed to hear a new sequence, signalling the end of the round. Participants were then given another minute to familiarise themselves with the range of movements they could make and ensure the volume and headphone fit were comfortable. Once they were satisfied, the experimenter would start the script. The experimenter remained in the room with the participants for the first 3 sequences (of the 32 warm-up trials), to ensure they had understood the procedure and the code was working. At the end of the last round, participants filled in an information sheet asking for their age, gender and degree of familiarity with music, before leaving.

## 2. Model: Latent Variables

### 2.1. Data

Let 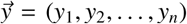 be a time series of onsets for one drum pattern. Let 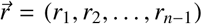 be the series of interval ratios, such that 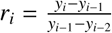.

### 2.2. Clustering datapoints into interval ratio categories

We assume that participants do not just perceive and represent the absolute datapoints (*r*_*i*_) veridically, but potentially reduce the variation among datapoints by assigning each to a rhythmic category. These categories are assumed to be learned in an unsupervised fashion by the learner. This category structure is modeled as a weighted mixture of *k* Gaussian distributions. Each category is defined by an unknown mean (*µ*_*k*_) and variance (σ_*k*_), and its weight in the mixture (ω_*k*_), where 0 ω_*k*_ 1 and Σ_*k*_ ω_*k*_ = 1. Let 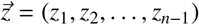 be a vector of category assignments indices such that we would write *z*_*i*_ = *k* if datapoint *r*_*i*_ has been assigned to rhythmic category *k*. When modelling a learner estimating these quantities, we assume a standard statistical model for Bayesian inference of an unknown mean and variance (the Normal-Inverse-Chi-Square model with uninformative prior parameters, see Gelman et al, 2006), and a standard statistical model for inference of the weights (weights 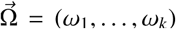 are modelled as a multinomial probability vector estimated under a uniform Dirichlet prior, see Gelman et al. 2006).

### 2.3. Chunking subseries into motifs

We wish to describe the structure in drum patterns as transitions between predictably structured, prototypical categories of subseries, or *motifs*. A motif is a kind (type), and we observe instances (tokens) of that motif in the data. Every subseries is an instance of a motif. Let 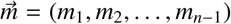 be notation that tells us which motif each datapoint has been assigned to. As before, *m*_*i*_ = *j* tells us that datapoint *r*_*i*_ has been judged to be part of an instance of motif *j*.

### 2.4. Boundary Variables

Allow the notation 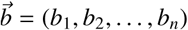 to denote the boundaries between subseries. Arbitrarily, we will keep track of the onset in a subseries (rather than the coda). If datapoint *r*_*i*_ is the first interval ratio of a motif, then *b*_*i*_ = 1, otherwise *b*_*i*_ = 0. By definition *b*_0_ = 1.

### 2.5. Motif internal Structure

We pursue one of the simplest possible formulations of motif categories: each category of motif is defined by a single prototype series of interval ratio categories. The probability that a newly observed subseries belongs to a particular category of motif depends on the subseries being an exact instance of the prototype sequence, or a close match.

## 3. Model: A sequential learning algorithm for the latent variables

Our focus is on approximating the representations of patterns that participants induce upon observing these patterns. As a result we choose to model individual participants using a lightweight, sequentially dependent posterior approximation algorithm for the latent variables defined by our model. We take McCauley and Christiansen’s (2012) CAPPUCINO model as inspiration for our algorithm. This model provides an online, sequential updating procedure that segments sequences of observed events (in their case words, in our case IOI ratios) and chunks these sequences into predictable subsequences by keeping track of transition probabilities between events. Here we describe our adaptation of this algorithm, and the additional variables it requires.

### 3.1. Transition Probabilities

The core assumption underpinning this algorithm is that individual use statistical information about the co-occurrence of interval ratio categories (*z*_*i*_) to make decisions about boundaries between motifs in patterns. Improbable sequences of interval ratio categories imply a boundary between motifs, because by definition a motif is a predictable sequence of interval ratio categories: transition probabilities between categories within a motif are likely to be high, whereas transition probabilities between the final category in a preceding motif and the onset category in a succeeding motifs are likely to be lower. Dips in transition probabilities provide cues to boundaries.

Following McCauley and Christiansen, our algorithm keeps track of *backwards* transition probabilities. For all *k* categories of interval ratio, we keep track of a vector of probabilities 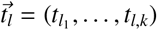 whose entries capture the probability that interval ratio category *l* is preceded by each other interval ratio category. Each vector *t*_*l*_ is a multinomial probability vector estimated under a uniform Dirichlet prior (unbiased Bayesian inference).

### 3.2. Creating and storing motifs

The algorithm described below assigns subseries to motifs based on the similarity between the subseries and a hypothesised motif. When a candidate subseries is an exact match for an existing motif which has been attested at least twice in previous patterns (this minimum count follows the CAPPUCINO model procedure), the model probabilistically assigns the new subseries to this motif category (we allow a small probability – 0.25 - that the model chooses instead to continue through the sequence before assigning a motif category). However, when a subseries of interval ratios is not an exact match for an existing motif, or if the model decides not to exploit the match, the model has the ability to create a new category of motifs entirely to account for the new datapoint. This decision process has a natural analogue statistical model in the Dirichlet Process, which is commonly formulated sequentially like our procedure. For any existing category of motifs, the model checks whether the motif matches the prototype motif, and assigns a likelihood of 1 to any matching categories. The prior for any motif category is the number of subseries already assigned to this category divided by a normalising constant *c* = *s* – 1 + α, where *s* is the total number of subseries assigned to all motifs, and α is a parameter of the model. During every decision, the model also entertains the possibility that a new motif category must be created to account for this subseries. The marginal likelihood of a new motif is linearly inversely proportional to its length (1 over the number of interval ratios in the subseries), and the prior probability of a new motif is α divided by the same normalising constant *c*.

This formulation corresponds closely to a sequential understanding of the Dirichlet process, in which short motifs are privileged, and a parameter α controls how happy the learner is to invent new motif categories. In our analyses, we set α = 10^−1^, which corresponds to the assumption that participants weakly prefer to re-use existing motif categories where possible. We run the model ten times on each participants’ data, and plot the average results in the main manuscript. This procedure is a close analogue of a particle filter, which is a commonly used technique in machine learning for sampling the posterior of a Dirichlet Process model.

### 3.3. Algorithm description

Here is procedure for running the algorithm to model an individual participant. We run this procedure only on the patterns observed (not produced), and only observed in the retest portion of the experiment (i.e. the final 32 patterns observed at each generation).

### initialization

1: construct an initial representation of interval ratio categories by categorizing, in order, every individual interval ratio in the initial sequence of 32 *test* patterns. For each individual datapoint, the categorisation procedure works by computing the posterior predictive distribution of each Gaussian interval ratio category, using this as the likelihood for the datapoint under consideration, and using the category mixture weight (ω_*k*_) as the category prior, then drawing a single sample from the posterior over categories.

2: Randomly sample initial transition probabilities from a uniform Dirichlet prior. During this sequence of categorisation decisions in step 1. of the initialization, update these transition probabilities sequentially by sampling the posterior, which is also Dirichlet thanks to the conjugacy of multinomial probability vector likelihoods and the Dirichlet prior.

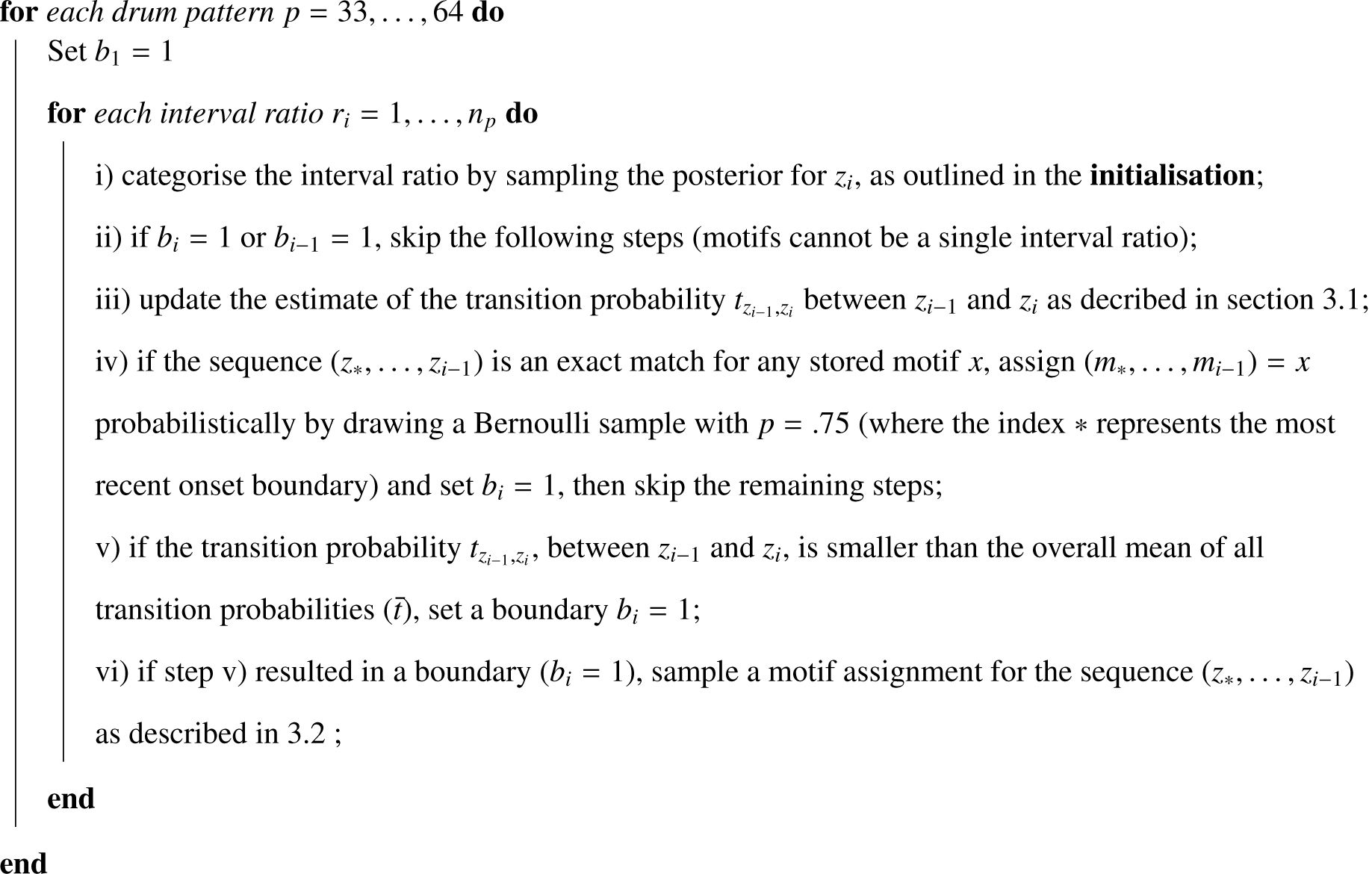

**Algorithm 1:** Psuedo-code for the learning algorithm implementing our model. Our algorithm is based closely on the CAPPUCINO model, and is a close analogue of a particle filter posterior sampling procedure.

## References

2. Honing, H., C. ten Cate, I. Peretz, et al. 2015. Without it no music: cognition, biology and evolution of musicality. Philosophical Transactions of the Royal Society of London B: Biological Sciences. 370: 20140088.

3. Fitch, W.T. 2017. Cultural evolution: Lab-cultured musical universals. Nature Human Behaviour. 1: 13

4. Jacoby, N. & J.H. McDermott. 2017. Integer Ratio Priors on Musical Rhythm Revealed Cross-culturally by Iterated Reproduction. Current Biology.

5. Ravignani, A., T. Delgado & S. Kirby. 2016. Musical evolution in the lab exhibits rhythmic universals. Nature Human Behaviour. 1: 0007.

6. Ravignani, A. & T. Verhoef. in review. Which melodic universals emerge from repeated signaling games? Artificial Life.

7. Lumaca, M. & G. Baggio. 2017. Cultural transmission and evolution of melodic structures in multi-generational signaling games. Artificial Life.

8. Le Bomin, S., G. Lecointre & E. Heyer. 2016. The Evolution of Musical Diversity: The Key Role of Vertical Transmission. PloS one. 11: e0151570.

9. Trehub, S.E. 2015. Cross-cultural convergence of musical features. Proceedings of the National Academy of Sciences. 112: 88098810.

10. Savage, P.E., S. Brown, E. Sakai, et al. 2015. Statistical universals reveal the structures and functions of human music. Proceedings of the National Academy of Sciences. 112: 8987-8992.

11. Rzeszutek, T., P.E. Savage & S. Brown. 2012. The structure of cross-cultural musical diversity. Proceedings of the Royal Society of London B: Biological Sciences. 279: 16061612.

12. Verhoef, T., S. Kirby & B. de Boer. 2014. Emergence of combinatorial structure and economy through iterated learning with continuous acoustic signals. Journal of Phonetics. 43: 5768.

13. Thompson, B., S. Kirby & K. Smith. 2016. Culture shapes the evolution of cognition. Proceedings of the National Academy of Sciences. 113: 45304535.

14. Miranda, E.R., S. Kirby & P. Todd. 2003. On computational models of the evolution of music: From the origins of musical taste to the emergence of grammars. Contemporary Music Review. 22: 91111.

15. Kirby, S., T. Griffiths & K. Smith. 2014. Iterated learning and the evolution of language. Current Opinion in Neurobiology. 28: 108114.

16. Kirby, S., M. Dowman & T.L. Griffiths. 2007. Innateness and culture in the evolution of language. Proceedings of the National Academy of Sciences. 104: 52415245.

17. Kirby, S., H. Cornish & K. Smith. 2008. Cumulative cultural evolution in the laboratory: An experimental approach to the origins of structure in human language. Proceedings of the National Academy of Sciences. 105: 1068110686.

18. Ravignani, A., H. Honing & S.A. Kotz. 2017. The evolution of rhythm cognition: Timing in music and speech. Frontiers in human neuroscience. 11.

19. Kirby, S. 2017. Culture and biology in the origins of linguistic structure. Psychonomic bulletin & review. 24: 118137.

20. Navarro, D.J., A. Perfors, A. Kary, et al. "When extremists win: On the behavior of iterated learning chains when priors are heterogeneous". In CogSci 2017.

21. Fehér, O., I. Ljubičić, K. Suzuki, et al. 2017. Statistical learning in songbirds: from self tutoring to song culture. Phil. Trans. R. Soc. B. 372: 20160053.

22. Griffiths, T.L. & M.L. Kalish. 2007. Language evolution by iterated learning with Bayesian agents. Cognitive science. 31: 441480.

23. Jamieson, R.K. & D. Mewhort. 2009. Applying an exemplar model to the artificial grammar task: Inferring grammaticality from similarity. The Quarterly Journal of Experimental Psychology. 62: 550575.

24. Ravignani, A. & G. Madison. in press. The paradox of isochrony in the evolution of human rhythm. Frontiers in psychology.

25. Levenshtein, V.I. 1966. "Binary codes capable of correcting deletions, insertions, and reversals". In Soviet physics doklady, Vol. 10: 707710.

26. Lee, C.y. & J. Glass. 2012. "A nonparametric Bayesian approach to acoustic model discovery". In Proceedings of the 50th Annual Meeting of the Association for Computational Linguistics: Long Papers Volume 1: 4049. Association for Computational Linguistics.

27. McCauley, S.M. & M.H. Christiansen. 2011. "Learning simple statistics for language comprehension and production: The CAPPUCCINO model". In Proceedings of the Cognitive Science Society, Vol. 33.

28. Motz, B.A., M.A. Erickson & W.P. Hetrick. 2013. To the beat of your own drum: Cortical regularization of non-integer ratio rhythms toward metrical patterns. Brain and cognition. 81: 329336.

29. Desain, P. & H. Honing. 2002. The formation of rhythmic categories and metric priming. Perception. 32: 341365.

30. Honing, H. & D. Deutsch. 2013. Structure and interpretation of rhythm in music. The psychology of music. 369404.

31. Lumaca, M. & G. Baggio. 2016. Brain potentials predict learning, transmission and modification of an artificial symbolic system. Social cognitive and affective neuroscience. 11: 19701979.

32. Lumaca, M., A. Ravignani & G. Baggio. in review. Music evolution in the laboratory: Cultural transmission meets neurophysiology. Frontiers in Neuroscience.

